# Spatially resolved gene expression and tumour-microenvironment interactions across malignant transformation in IDH-mutant glioma

**DOI:** 10.64898/2026.09.08.749421

**Authors:** Alyona Ivanova, Shamini Ayyadhury, David G Munoz, Megan Hopkins, Farzaneh Aboualizadeh, Melanie Spears, Megan Wu, Trevor Pugh, Roel G W Verhaak, Daniel P Cahill, Sunit Das

**Affiliations:** The Arthur and Sonia Labatt Brain Tumour Research Center, The Hospital for Sick Children, Toronto, Ontario, Canada; Institute of Medical Sciences, University of Toronto, Toronto, Ontario, Canada; Princess Margaret Cancer Centre, University Health Network, Toronto, Ontario, Canada; Department of Laboratory Medicine, St. Michael’s Hospital, University of Toronto, Toronto, Ontario, Canada; Department of Laboratory Medicine and Pathobiology, University of Toronto, Toronto, Ontario, Canada; Ontario Institute for Cancer Research, Toronto, Ontario, Canada; Department of Medical Biophysics, University of Toronto, Toronto, Ontario, Canada; Harvey and Kate Cushing Professor of Neurosurgery, Yale University, New Haven, CT, the USA; Department of Neurosurgery, Massachusetts General Hospital and Harvard Medical School, Boston, MA, USA; Division of Neurosurgery, University of Toronto, Toronto, Canada

**Keywords:** IDH-mutant glioma, astrocytoma, migration, invasiveness, spatial transcriptomics

## Abstract

Isocitrate dehydrogenase (IDH)-mutant gliomas represent most lower-grade diffuse gliomas in young adults. Although IDH-mutant gliomas initially behave indolently relative to IDH-wildtype counterparts, they undergo inevitable malignant transformation. The mechanisms driving this progression remain poorly defined. We analyzed a unique longitudinal cohort of two patients with IDH-mutant diffuse gliomas, each with matched WHO grade 2, 3, and 4 tumours. Spatial transcriptomics (NanoString GeoMx and 10X Visium), combined with differential expression, spatially variable gene analysis, and gene ontology enrichment, were used to define transcriptional programs underlying progression. Unexpectedly, across both patients, higher-grade tumours demonstrated reduced proliferation at the infiltrative margin as measured by Ki67. Spatial transcriptomic analysis identified an enrichment of invasion-associated genes associated with increasing tumour grade. In an independent bulk RNA-seq cohort, expression of this spatially derived program did not outperform grade-only PFS or OS analysis. These observations underscore the need to develop new metrics that account for spatial evolution that occurs across progression, but escapes detection in bulk analyses and provide the rationale for prioritizing migration-, cytoskeletal- and stress-associated programs for further investigation.

**Importance of the Study:** Gliomas with IDH mutations exhibit a more favorable prognosis than IDH–wildtype gliomas but inevitably undergo further malignant transformation over time. Understanding the molecular trajectory of malignant transformation will be critical to determine the indications for IDH inhibitor therapy in patients with these tumours. Identifying actionable molecular programs will be key to generate precision therapeutic strategies for patients with IDH-mutant tumours.

## INTRODUCTION

Gliomas with mutations in the isocitrate dehydrogenase (IDH) genes constitute the majority of lower-grade diffuse gliomas seen in young adults. IDH mutations produce the oncometabolite D-2 hydroxy-glutarate (D-2HG) at the expense of Alpha-Ketoglutarate (α-KG). Elevated D-2HG competitively inhibits α-KG–dependent dioxygenases, while α-KG depletion further disrupts epigenetic regulation, contributes to pathological self-renewal of stem-like progenitor cells, impairs DNA repair, increases immunosuppression, together promoting impaired differentiation and gliomagenesis.^1–4^ Lineage and differentiation based studies indicate that IDH mutant gliomas are characterized by a blocked oligodendrocyte lineage differentiation program, which may shape tumour behavior and response to therapy.^5^ The development and clinical integration of the IDH inhibitor, vorasidenib, and the hope that these agents may modify the biological effects of IDH mutation in these tumours, has highlighted the importance of understanding how IDH mutations shape the molecular trajectory of glioma progression.

Although IDH-mutant gliomas exhibit a more favorable prognosis than IDH-wildtype tumours, they inevitably undergo malignant transformation over time^6–8^. Current therapies, including surgical resection, radiotherapy, chemotherapy and targeted therapy, have been shown to improve upon the natural history of these cancers but fail to prevent tumour progression and death^9–10^. Emerging studies to map the evolutionary trajectories of IDH mutant gliomas have revealed that while early stage tumours share common molecular features, the patterns of progression diverge significantly, depending on subtype and grade, resulting in heterogeneity that is difficult to mitigate and control.^11–12^

Tumour cells dynamically oscillate between migration and proliferation states depending on environmental cues, contributing to therapy resistance and recurrence. Hypoxia, nutrient deprivation, and mechanical stress often trigger the “Go” phenotype, promoting motility and invasion, while growth factors, nutrient abundance, and cell–cell contact promote the “Grow” phenotype, enhancing cellular proliferation.^13–14^ In astrocytic tumours, highly migratory cells at the tumour margins are often quiescent or slowly dividing, making them less sensitive to conventional therapies that target rapidly proliferating cells.^13–14^ As astrocytomas progress to higher grades, they undergo mesenchymal (MES) subtype switching, also known as proneural-mesenchymal transition (PMT). This switch is strongly associated with acquiring a more invasive, less proliferative phenotype and is reflected by genetic changes.^15^

While we have a well-developed understanding of the histological and molecular characteristics of IDH-mutant tumours progression, which has resulted in the current version of molecular classification of gliomas, the mechanisms that drive this process remain incompletely defined, limiting the development of effective timepoint-specific therapeutic strategies. Our study aims to leverage a rare longitudinal spatial cohort of patient-matched gliomas (n=2) spanning across the tumour grade spectrum to identify candidate expression patterns for subsequent validation in larger longitudinal cohorts.

## RESULTS

We identified two patients with IDH-mutant diffuse infiltrative glioma for whom archived tissue representing progression from WHO grade 2, 3 and 4 disease, were available (Supplementary Table 1, **Figure 1**).

**Figure 1.**
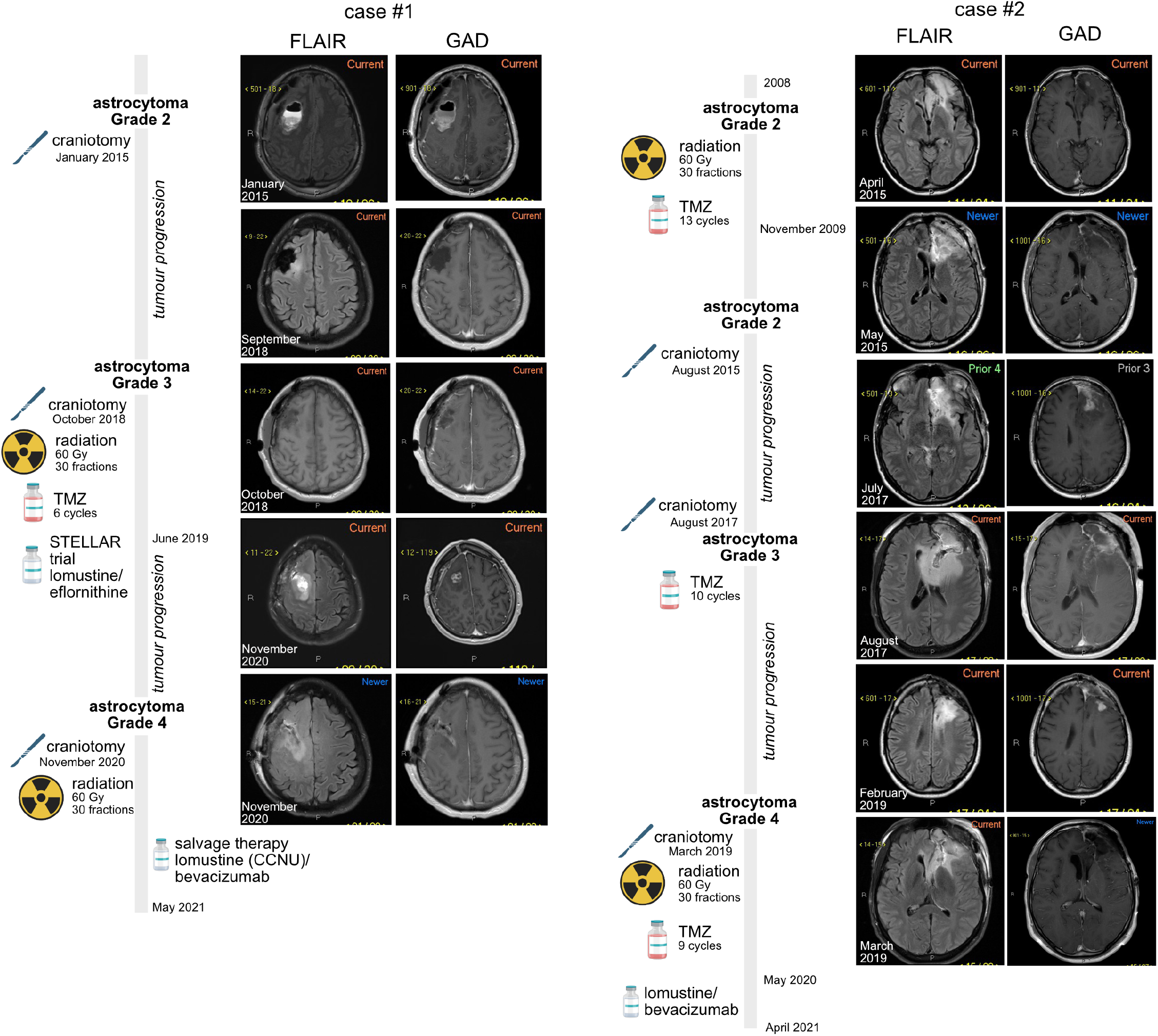
Representative MRI images (FLAIR and GAD-enhanced T1-weighted sequences) illustrating disease progression over time, alongside the patients’ clinical course and treatment history. Tissue for spatial profiling was collected at craniotomy.

### Clinical history

#### CASE 1

This patient underwent craniotomy for a non-enhancing, FLAIR hyperintense left frontal WHO grade 2 astrocytoma, IDH-mutant, performed after his presentation with seizure, in January 2015. He was followed with active surveillance until October 2018, when progression of non-enhancing disease prompted a right frontal craniotomy, with pathology confirming transformation to grade 3 astrocytoma, IDH-mutant. The patient underwent radiation (60 Gy in 30 fractions) with concurrent and six adjuvant cycles of temozolomide (TMZ), completed in June 2019. Serial surveillance showed evidence of slow expansion of FLAIR-hyperintense signal adjacent to the resection cavity. He was enrolled in the STELLAR trial, receiving one cycle of lomustine and eflornithine, but returned to the operating room in November 2020 following an MRI showing new tumour enhancement. Repeat craniotomy revealed transformation to grade 4 astrocytoma, IDH-mutant, with MGMT promoter methylation. On WES analysis, the patient presented with a treatment-induced hypermutated genetic signature ^30^, including *IDH1* mutation (R132H), *ATRX* loss, *FUBP1* splice mutation, *POLE1* missense (F695I) and splice mutations, 5’ *TERT* promoter mutation, missense mutations in *SMARCA4* and *TP53* (**Figure 2**). Other alterations included *ALK* (I1461V) and *MET* missense, *CCND1* and *GNAQ, NOTCH1, NOTCH3, PI3KCA, STK11* splice mutations, *DICER1* loss. The patient underwent re-irradiation, completed in January 2021, and received two cycles of salvage therapy with lomustine (CCNU) and bevacizumab, terminated at disease progression. The patient expired in May 2021.

**Figure 2.**
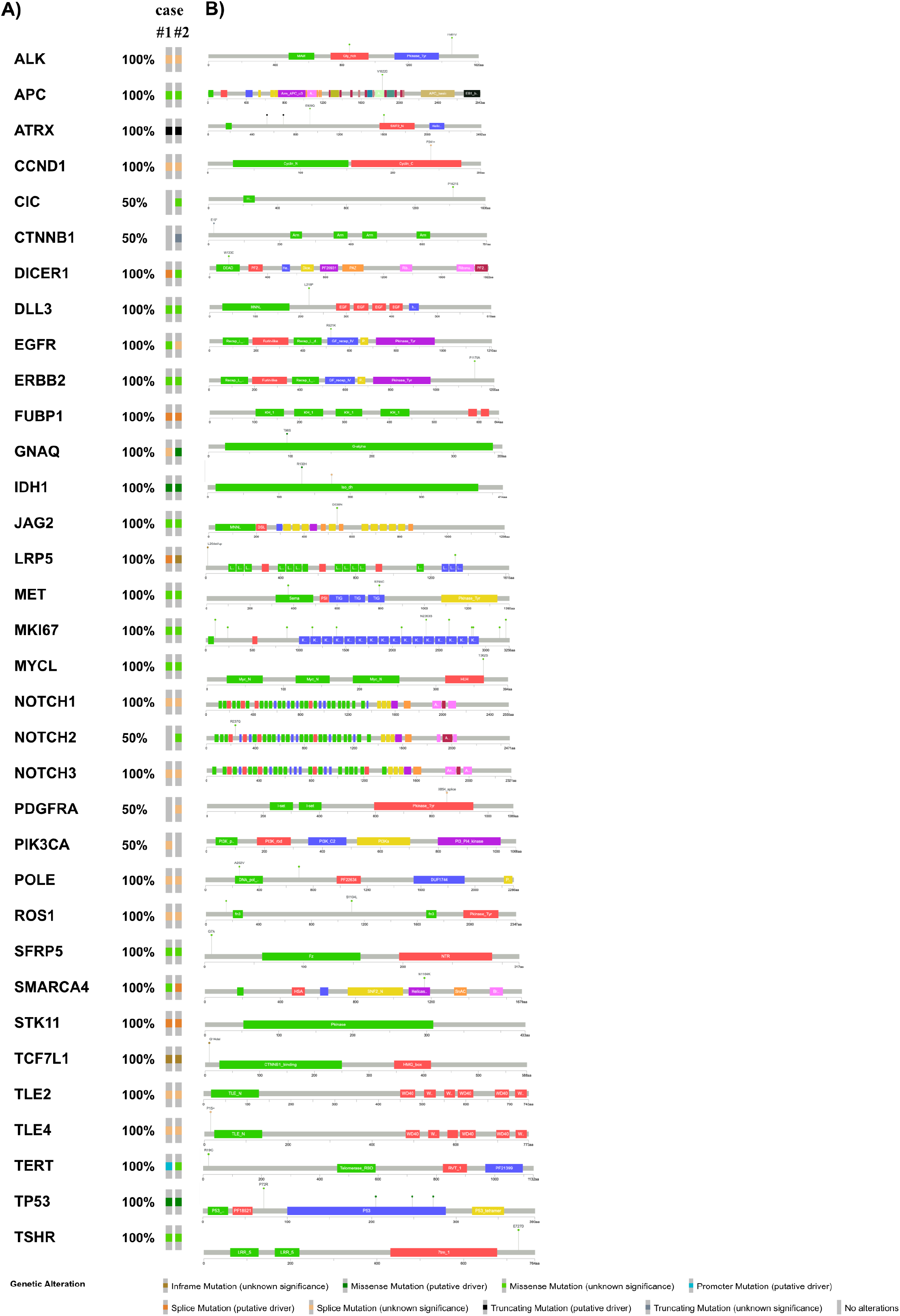
Genomic landscape of IDH-mutant astrocytomas. (A) Oncoplot demonstrating the distribution of somatic alterations across both cases. Columns represent individual tumour samples and rows represent altered genes. Mutation classes are color-coded by alteration type (missense, nonsense, splice-site, amplification, or structural variant). Percentages represent frequency of alteration in patient cohort. (B) Lollipop plots of selected genes showing the positional distribution of coding variants relative to functional protein domains. Splice-site and missense variants are shown according to genomic position.

#### CASE 2

This patient was initially diagnosed with a WHO grade 2 glial tumour (IDH-mutant) on biopsy performed on his presentation with seizure in 2008. The patient was treated with TMZ chemoradiation (60 Gy/30 fractions) followed by 13 cycles of adjuvant TMZ, completed November 2009. He returned to surgery in August 2015, prompted by radiographic findings of slow, non-enhancing tumour progression. Pathology demonstrated a WHO grade 2 astrocytoma, IDH-mutant with ATRX loss. Surveillance imaging showed new tumour enhancement in August 2017. He returned to surgery. Pathology revealed a WHO grade 3 astrocytoma, IDH-mutant, with ATRX loss and retained 1p and 19q. The patient received 10 additional cycles of adjuvant TMZ, but returned to surgery in March 2019, when surveillance imaging showed evidence of new tumour enhancement. Pathology revealed progression to WHO grade 4 astrocytoma, IDH-mutant. On WES analysis, the patient presented with a treatment-induced hypermutated genetic signature ^30^(**Figure 2**), including *IDH1* mutation (R132H), *ATRX* loss, *FUBP1* and *SMARCA4* splice mutations, *POLE1* missense (A252V), *TP53* missense, *TERT* missense (R19C). Other oncogenic mutations identified were *ALK* (I1461V), *DICER1, MET, NOTCH2* missense, *EGFR, GNAQ, NOTCH1* and *NOTCH3, PDGFRA, STK11* splice mutations, *CTNNB1* nonsense mutation. Patient underwent repeat radiation (60 Gy/30 fractions) with concurrent and 9 cycles of adjuvant TMZ, completed May 2020. He was subsequently treated at disease progression with three cycles of lomustine and bevacizumab, initiated in July 2020 and discontinued at progression after three cycles. He expired in April 2021.

### Defining spatial gene expression across progressive timepoints of IDH-mutant glioma transformation

Pathologic review of the cases showed progressive diagnostic features of malignant transformation, with grade 2 samples displaying an infiltrative glial tumour with considerable nuclear atypia, and grade 4 demonstrating a high degree of anaplasia with marked nuclear pleomorphism and hyperchromatism (**Supplementary Table 1**). Tissue sections for spatial analysis were chosen by a neuropathologist based on histological features. Specifically, we identified the infiltrative edge area of the tumour (invasive front) as the transition between infiltrating tumour cells and adjacent non-neoplastic brain parenchyma where individual tumour cells extend beyond the main tumour mass.

#### CASE 1

Using immunostaining for GFAP and Ki67 on the NanoString GeoMx platform, we spatially resolved tumour regions (GFAP+) and quantified proliferation rate based on Ki67 protein expression (**Figure 3A-C**, **Table 1**). Grade 4 tumours had lower Ki67 index compared to grades 2 and 3, indicating a less proliferative phenotype with disease progression. Differential expression analysis identified enrichment of migration-related genes in ROIs from grade 4, compared to grade 2 and 3, tumours (**Table 2**) including *TTYH2*, *DNAJB2*, *TUBB4A*, *MYH2* (**Figure 3D-E**). *TTYH2*, *DNAJB2* and *MYH2* expression was also associated with mesenchymal gene enrichment: *ZEB1* and *SOX4* (**Figure 3E**). On gene set enrichment analysis, pathways involved in nervous system development, axon guidance, and cell adhesion were associated with grade 4 tumours (**Figure 3F**).

**Figure 3.**
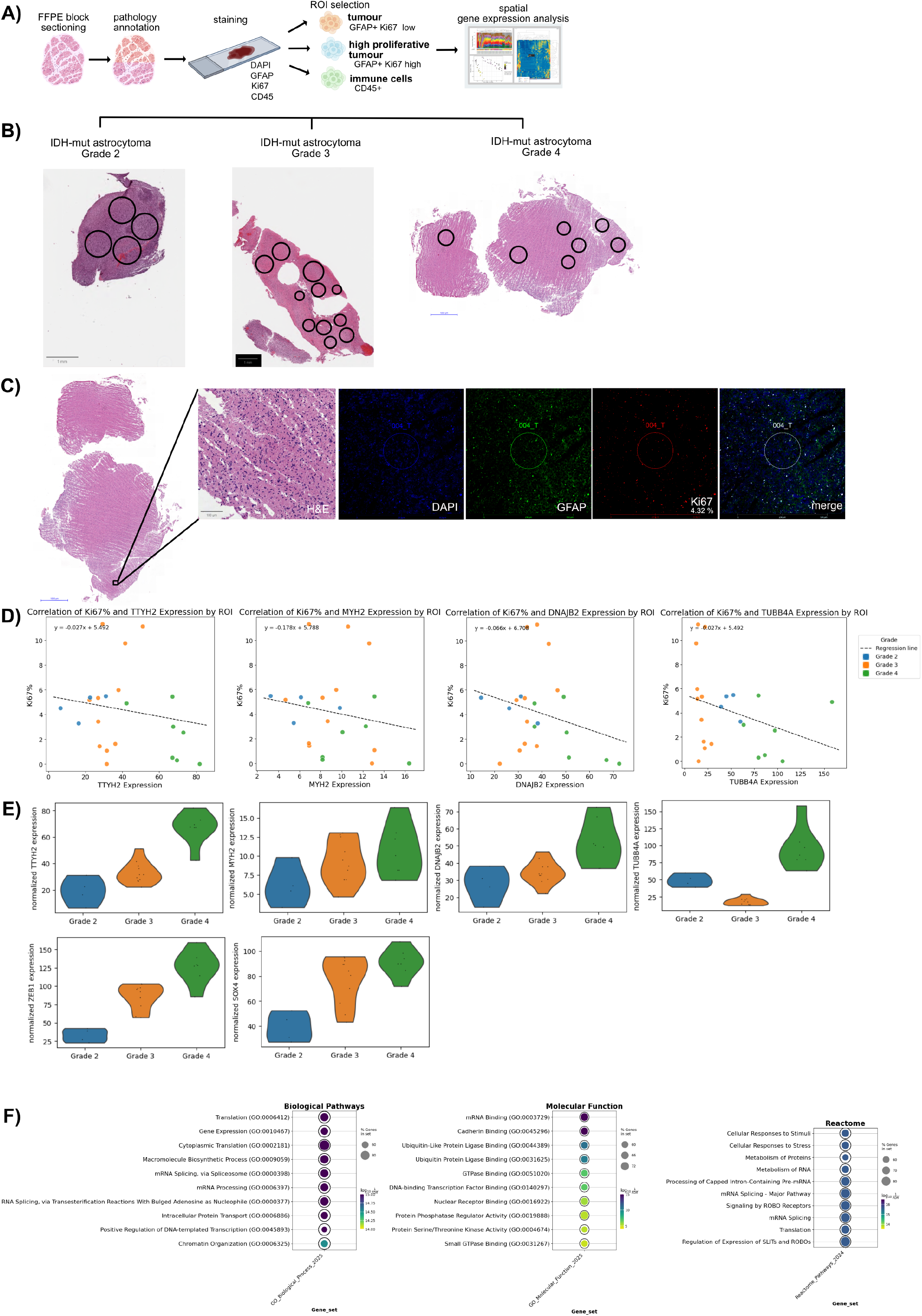
CASE 1 (A) GeoMx workflow. (B) H&E of longitudinal IDH-mut samples submitted for GeoMx profiling. Scale 1mm. (C) Representative IDH-mutant grade 4 tumour depicting invasive low-proliferative cells. Stained for H&E, DAPI, and markers GFAP and Ki67. Scale 1mm, 100µm. (D) Correlation plots of normalized gene expression in each ROI versus Ki67 protein expression for each tumour grade. Linear regression demonstrates an inverse relationship between proliferation index and invasion-associated transcriptional programs across disease progression. (E) Violin plots depicting increased expression of invasion-associated genes and mesenchymal state genes across progressive grades. The dots correspond to normalized gene expression in each individual ROI. (F) GO Biological Processes, Molecular Function, and Reactome enrichment analysis of genes upregulated in grade 4 tumour.

**Table 1.**
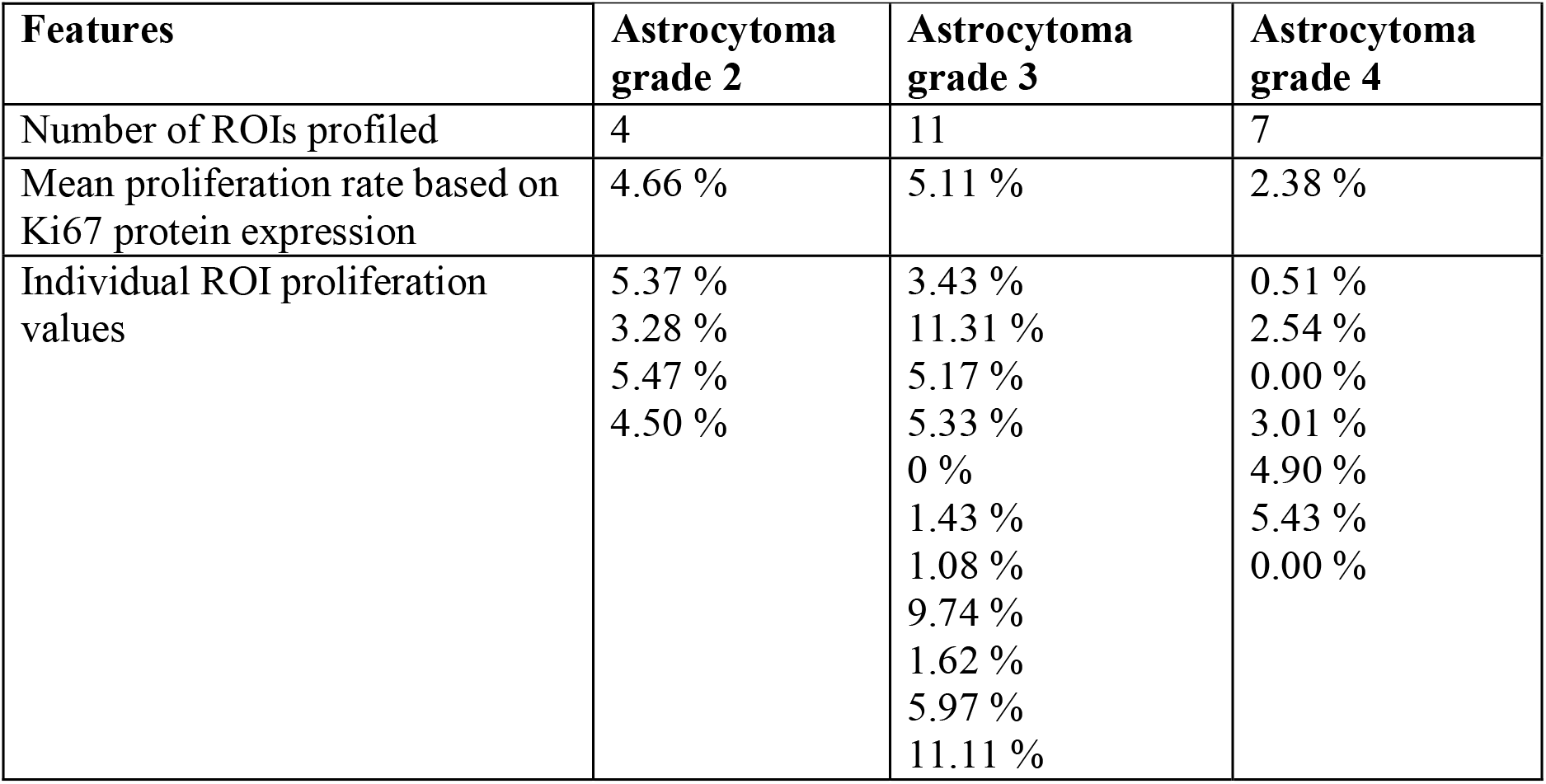
CASE 1: GeoMx key features. Table 1 lists the number of ROIs profiled for each tumour section (grade 2, grade 3, grade 4) of CASE 1, individual ROI proliferation values based on percentage of Ki67 positive cells, and the mean Ki67 protein expression across ROIs for each tumour section.

**Table 2.** Migration-associated genes identified by preliminary spatial profiling of IDH-mutant glioma. Candidate genes were selected based on their biological relevance to cellular migration, invasion, cytoskeletal organization, or tumour–microenvironment interactions and their increased expression in grade 4 tumour regions relative to lower-grade tumour regions in the preliminary spatial profiling dataset. Log2 fold change (L2FC) values represent the relative expression of each gene in grade 4 versus lower-grade tumour regions for Case 1 and Case 2, respectively. Positive L2FC values indicate higher expression in grade 4 regions. “n/a” indicates that the gene was not detected. Because the preliminary dataset contains limited biological replication, these comparisons are considered hypothesis-generating and are not interpreted as independent patient-level statistical evidence of differential expression.

| Gene name | Function | grade 4 vs lower grades CASE 1 | grade 4 vs lower grades CASE 2 |
| --- | --- | --- | --- |
|  |  | L2FC | L2FC |
| ***TTYH2*** | Critical for invasion and migration, associated with formation of tumour microtubules in glioma | 1.25 | 1.52 |
| ***DNAJB2*** | Co-chaperone crucial for the proper migration of developing neural precursor cells | 0.72 | 1.96 |
| ***TUBB4A*** | Forms microtubules necessary for neuronal migration, protects the nucleus from damage during cell migration | 1.88 | n/a |
| ***MYH2*** | Motor protein that generates the mechanical force during cell migration by interacting with actin filaments to contract and retract the trailing edge of the cell, and to facilitate protrusions at the leading edge | 0.45 | n/a |
| ***EDIL3*** | Enhances cell migration, especially in angiogenesis and cancer, by promoting cell movement and invasion through interactions with integrins | 1.57 | 2.64 |
| ***SRCIN1*** | activates the Wnt/β-catenin signaling pathway and promotes migration | 1.43 | 4.98 |

#### CASE 2

In contrast to GeoMx, Visium does not require manual selection of ROIs but instead allows capture of unbiased, whole-mount data. For each progression timepoint (grade 2, 3 and 4), we identified tissue sections with heterogeneous morphology to capture continuous gene expression at the non-proliferating invasive tumour margins (**Supplementary Table 1**, **Figure 4A**). Leiden clustering (Visium spots) of integrated gene expression data across disease timepoints allowed to identify genetically similar clusters of cells (**Figure 4B**). We focused on the same set of invasive genes identified in Case 1.

**Figure 4.**
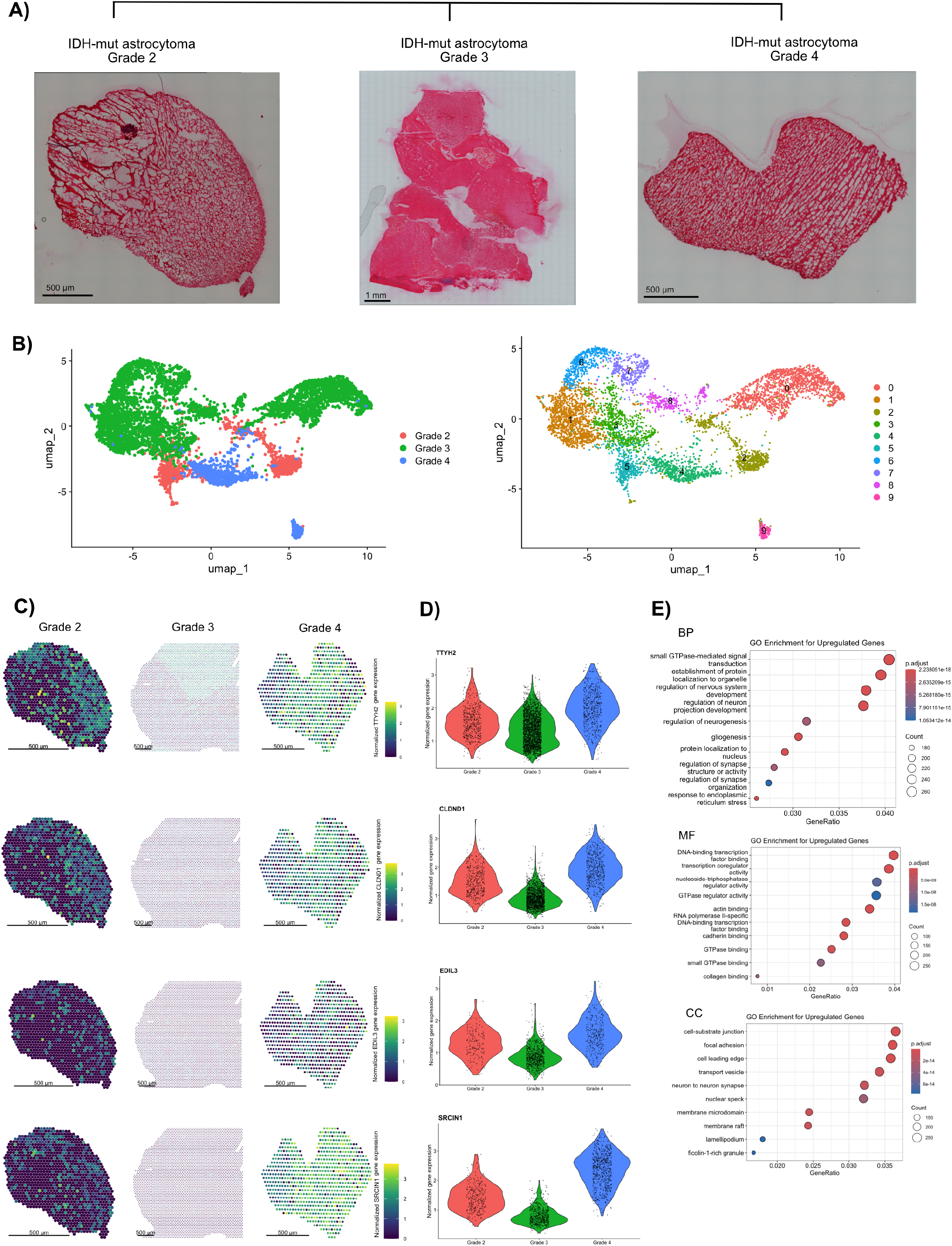
CASE 2 (A) H&E of longitudinal IDH-mut samples submitted for Visium profiling acquired at 10X magnification. (B) UMAP plots depicting histological grade (left) and spatial clusters following joint leiden clustering (right). (C) Spatial maps showing expression of invasiveness-associated genes in tumours of different grades. Scale bar 500 µm. (D) Violin plots showing gene expression of invasiveness-associated genes in tumours of different grades. The dots correspond to spot-level normalized gene expression. (E) Biological Processes, Molecular Function, and Cellular Component GO enrichment analysis of genes upregulated in grade 4 tumour.

**Figure 5.**
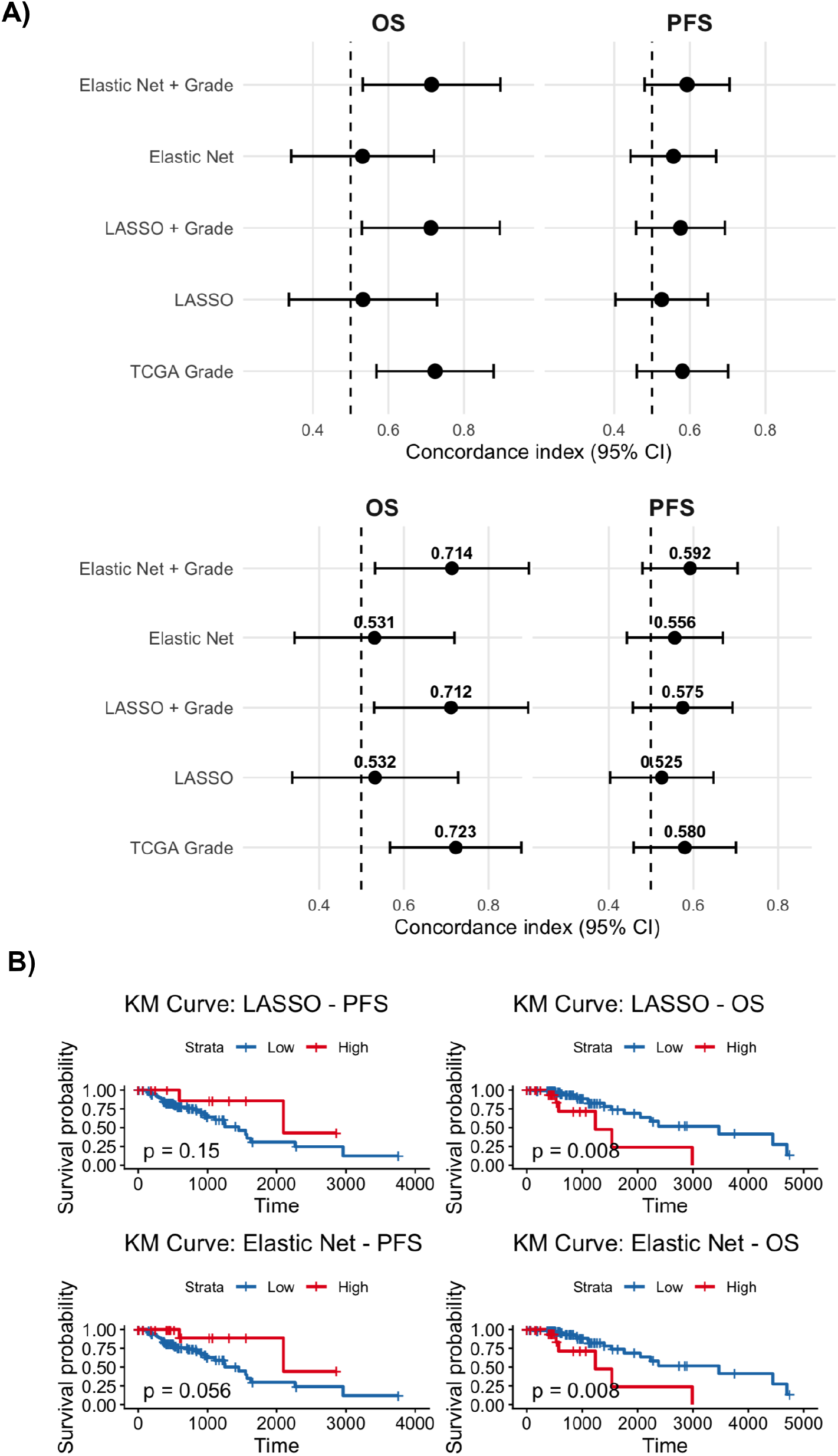
(A) Test-set discrimination of regularized prognostic model inputs in longitudinal IDH-mutant patient samples from The Cancer Genome Atlas (n=446)^24^. Concordance indices (C-indices) with 95% confidence intervals are shown for LASSO, elastic-net, and TCGA tumour grade models for OS and PFS in the independent test cohort. Models incorporating TCGA tumour grade are shown alongside invasiveness gene-score-only models. The number is the C-index, while the horizontal line is its 95% CI. The dashed vertical line represents a C-index of 0.50, corresponding to chance-level discrimination. Confidence intervals were calculated as C-index ± 1.96 × standard error. (B) Kaplan–Meier curves for OS and PFS in the held-out test cohort stratified into high- and low-risk groups according to the regularized gene scores derived from LASSO and elastic-net Cox models. P values were calculated using log-rank tests. Risk groups were defined using the maximally selected rank statistic.

As in Case 1, proliferation decreased with increasing tumour grade, as characterized by *MKI67* expression. *TTYH2* was expressed across all tumour grades (**Figure 4C-D**) but showed slight enrichment in expression in grade 4 disease. On Differential Gene Expression (DEG) analysis, we again found evidence of enriched expression of tumour invasiveness genes with advancing grade (**Figure 4C-D**, **Table 2**, see also **Supplementary Table 2**), including cadherins (*CDH9* and *CDH12*) and other adhesion molecules (*NCAM2*, *CNTN2*), ephrins (*EPHA7* and *EPHA10*), semaphorins (*SEMA4D*), coronins (*CORO6*), RAS-family regulatory proteins (*RASAL1*), and RNA binding proteins (*CELF4*).

In addition to DEG analysis, Visium also enables spatially variable gene (SVG) analysis, which incorporates spatial coordinates to uncover patterns of gene expression that are organized across the tissue architecture, and is thus able to uncover localized molecular programs, such as cell adhesion and migration, that contribute to spatial heterogeneity. SVG analysis identified enrichment of multiple DEGs involved in cell migration in the invasive tumour margin, e.g. *TTYH2*, *CLDND1*, *EDIL3*, *SEMA4D*, *CORO6*, *CNTN2*, *CELF4*, and *SRCIN1*) (**Figure 4C-D**). *L1CAM*, *EPHB6*, *PACSIN1* were also enriched at the tumour invasive margin in grade 4 tumours.

Gene Ontology (GO) enrichment analysis of genes upregulated in grade 4 tumours revealed activation of biological processes related to neural development and cellular structural organization, specifically, regulation of neuron projection development, regulation of nervous system development, and gliogenesis, including *SRCIN1*, *CNTN2*, *VIM*, *SEMA4D*, and *DNAJB2* (**Figure 4E**).^16–23^ Enrichment was observed for cadherin binding and actin binding, as well as for cellular component terms including focal adhesion, cell leading edge, and lamellipodium (**Figure 4E**). Notably, *VIM*, *ACTB*, and *CD44* were strongly associated with these cellular components, supporting a role for cytoskeletal dynamics and adhesion structures in this case of IDH-mutant glioma malignant transformation.

### Prognostic significance of an invasiveness-related gene signature in relation to tumour grade

To investigate the prognostic relevance of genes identified from spatial transcriptomic analysis, 22 candidate genes derived from differential expression analysis (**Supplementary Table 3**) were evaluated using LASSO-penalized Cox regression in 446 IDH-mutant astrocytoma patients (grades 2-4) from the TCGA Pan-Cancer Atlas.^24^ Because TCGA RNA-seq data are derived from bulk tumour specimens, this analysis does not directly validate the spatial localization of the identified program, but rather provides an independent assessment of whether expression of genes comprising the spatially derived program is associated with clinical outcome. For OS, LASSO selected *TTYH2, CNTN2, PLEKHG3* and *DAAM2* as the final signature. In the independent test cohort, the LASSO-derived score was not associated with OS (HR = 1.00, 95% CI 0.43–2.31, p = 0.996; C-index = 0.532). In comparison, TCGA tumour grade demonstrated greater discrimination (C-index = 0.723), with Grade 4 associated with increased risk of death relative to Grade 2 (HR = 17.28, 95% CI 4.08–73.29). Addition of the LASSO score to TCGA grade did not significantly improve model fit (likelihood-ratio test, p = 0.366) or discrimination (C-index = 0.712 vs. 0.723 for grade alone). In the adjusted model, the LASSO score was not independently associated with OS (HR = 1.62, 95% CI 0.57–4.62, p = 0.36).

For PFS, LASSO selected *TTYH2, CNTN2, DAAM2, CD44, SRCIN1, SEMA4D, CDH9*, and *RASAL1.* The LASSO-derived score was not associated with PFS in the independent test cohort (HR = 1.18, 95% CI 0.43–3.18, C-index = 0.525). TCGA grade showed greater discrimination (C-index = 0.580), with Grade 4 associated with increased risk of progression relative to Grade 2 (HR = 3.45, 95% CI 1.09–10.91). Addition of the LASSO score to TCGA grade did not improve model fit (likelihood-ratio test, p = 0.868) or discrimination (C-index = 0.575 vs. 0.580 for grade alone), and the adjusted LASSO score was not independently associated with PFS (HR = 1.04, 95% CI 0.31–3.46).

As a sensitivity analysis, Elastic Net Cox regression yielded similar discrimination and did not provide evidence of additional prognostic information beyond TCGA grade. The Elastic Net retained *TTYH2* and *DAAM2* for OS and *TTYH2, DNAJB2, MYH2, CNTN2, DAAM2, PLEKHG3, SRCIN1, CRYAB, CD44, CDH9, RASAL1, EPHA10, CDH12* and *SEMA4D* for PFS. In the independent test set, Elastic Net scores were not associated with OS (HR = 0.94, 95% CI 0.30– 2.96, p = 0.916; C-index = 0.531) or PFS (HR = 1.41, 95% CI 0.55–3.61, p = 0.478; C-index = 0.56). After adjustment for TCGA grade, Elastic Net scores remained non-significant for OS (HR = 1.73, 95% CI 0.53–5.65; C-index = 0.714) and PFS (HR = 1.27, 95% CI 0.45–3.57; C-index = 0.592), and neither model significantly improved model fit over grade alone (OS, likelihood-ratio test p = 0.378; PFS, p = 0.657). Together, these findings indicate that the spatial transcriptomic gene program did not provide independent prognostic information beyond TCGA tumour grade for either OS or PFS in the independent cohort.

## DISCUSSION

In this study, we performed longitudinal spatial analysis of two IDH1-mutant gliomas through the full trajectory of malignant transformation.

Genetically, grade 4 tumours of both patients demonstrated similar molecular profiles, likely therapy-induced, including *IDH1* (R132H), *ATRX* loss, *TP53* missense mutations, and additional alterations involving *POLE*, *SMARCA4*, and splice-site variants in other cancer-associated genes. The *POLE* F695I variant is a somatic exonuclease-domain mutation predicted to impair DNA proofreading, resulting in an ultramutated phenotype. Pathogenic *POLE* alterations have been described in high-grade gliomas and other solid tumours, particularly in younger patients, where they are associated with markedly elevated tumour mutation burden (TMB) and distinctive mutational signatures. Defective polymerase proofreading may enhance neoantigen generation and has been linked to improved responsiveness to immune checkpoint blockade in selected cancers.^34–36^ *TP53* mutation is a canonical early driver event in IDH-mutant astrocytoma and frequently co-occurs with ATRX loss during gliomagenesis. Hotspot mutations such as R273C may confer gain-of-function properties that promote tumour progression, genomic instability, invasion, and resistance to apoptosis beyond simple loss of tumour suppressor activity.^37–39^ Loss of *DICER1* may further contribute to aggressive behaviour through disruption of microRNA biogenesis, leading to widespread post-transcriptional dysregulation. Impaired *DICER1* function has also been associated with accumulation of R-loops, replication stress, and chromosomal instability.^40^ *EGFR* abnormalities, although less common in IDH-mutant astrocytomas than in IDH-wildtype glioblastoma, have been associated with higher grade transformation, aggressive clinical behaviour, and inferior progression-free and overall survival.^41–42^ Similarly, *MET* alterations have been implicated in glioma invasion, stemness, and progression to higher-grade disease.^43^ The Notch signaling pathway is more commonly altered in oligodendrogliomas rather than IDH-mutant astrocytomas, where it regulates cell plasticity, maintains tumor-propagating cells, and contributes to disease progression, often through copy number losses of key Notch genes. Inactivating alterations in Notch genes are frequent, and their presence is associated with shorter PFS. In the cases described, Notch mutations are likely to be an example of passenger mutations or hypermutated events versus true hotspot ECD Notch inactivating mutations. Additionally, *PDGFRA* alterations may drive constitutive receptor signaling and activate downstream PI3K/AKT/mTOR and RAS/MAPK pathways, promoting proliferation and survival. Concurrent *PIK3CA* mutations may further amplify this oncogenic signaling axis.^44–45^ Finally, some studies suggest that *TERT* promoter mutations in IDH-mutant astrocytomas may identify biologically aggressive subsets with less favourable outcomes, although their prognostic significance remains context dependent.^46–47^ The non-canonical *TERT* (R19C) mutation is likely attributed to the therapy-induced hypermutation. Overall, defects in DNA repair (POLE), cell-cycle regulation (*TP53, CCND1*), receptor tyrosine kinase signaling (*EGFR, MET, PDGFRA, PIK3CA*), chromatin remodeling (*SMARCA4*), suggests mechanisms driving malignant progression. Due to tissue availability and quality, WES could not have been performed retrospectively in grade 2 samples to understand the genetic origin of the tumors prior to therapy.

While WES detects DNA mutations and copy number variations to identify genetic drivers and treatment-induced genetic changes, ST measures RNA levels in tissue, retaining spatial location. Spatial transcriptomic analysis suggested a shift from proliferative toward migratory phenotypes with advancing grade. Our preliminary spatial profiling identified a set of candidate genes, including *TTYH2, DNAJB2, MYH2* and *TUBB4A*, whose expression patterns were enriched in grade 4 relative to lower-grade tumours. These observations provided the rationale for prioritizing migration-, cytoskeletal- and stress-associated programs for further investigation. However, because these preliminary analyses were generated from a limited number of patients and contain multiple spatial observations per specimen, we will not interpret spot-level adjusted p-values as evidence of grade-level statistical significance. Our findings align with the canonical “Go-or-Grow” model, in which tumour cells dynamically alternate between proliferative and invasive states, depending on environmental cues,^13–14^ and astrocytic cells with a lower proliferation capacity contribute to more invasive phenotype. In our analysis, we identified the enrichment of DEGs and SVGs implicated in processes that drive local invasion and integration into neural circuits, such as *TTYH2*, *DAAM2*, *CDH9*, *CD44*, *CNTN2*, *SEMA4D* and *SRCIN1*.^16–19^ The deregulation of these programs in higher-risk groups suggests that enhanced migratory and synaptic-like signaling activity correlates with worse outcomes. These findings suggest a parallel between evolution in IDH-mutant gliomas and IDH-wild type glioblastomas, in which tumour cell-glial and tumour cell-neuronal interactions have been found to be critical to tumour progression and invasion.^16–20^

In this study, we used 2 different spatial platforms: GeoMx and Visium. Visium provides near-cellular resolution and unbiased transcriptome-wide coverage, enabling detailed mapping of transcriptional gradients within tissue. In contrast, GeoMx DSP allows for targeted high-plex profiling across histologically defined regions of interest, accommodating a wider dynamic range of tissue types and preservation states. Both platforms remain limited by their spatial resolution, as individual spots or regions often capture transcripts from multiple cells, challenging precise cell-type identification. To mitigate these limitations, scRNA-seq can serve as a complementary approach, enabling finer cell-type decomposition within spatial spots. Nevertheless, the small number of cells captured per spot in spatial transcriptomics data (considerably lower than in bulk RNA-seq) can introduce noise and confound deconvolution analyses, as demonstrated in several benchmark studies.^27–29^

A key limitation of the present study is that the derived molecular signature was generated from only two spatially profiled cases sampled across three longitudinal timepoints. Concordance of the gene set between cases is confounded by patient, lineage, tissue preservation (FFPE vs. fresh-frozen), and probe panel. Moreover, grade 4 samples are also the most heavily irradiated/TMZ-treated, so grade-associated transcription cannot be separated from therapy effects. The prognostic relevance of this signature did not show significant associations with PFS or OS in a larger cohort of 446 patients with IDH-mutant astrocytoma. However, the cohort remains limited in size and bulk sequencing data may not capture the full spatial biological heterogeneity of these tumors. Accordingly, future studies should prioritize expansion of the spatially resolved cohort and identification of molecular programs that demonstrate robust and reproducible associations with clinical outcomes across multiple independent datasets. Future studies should focus on functional validation of candidate invasive genes to determine whether they causally promote malignant progression and reduce dependency on canonical IDH-mutant signaling. Such findings may help identify mechanisms of resistance to IDH inhibitor therapy and inform future combination treatment strategies.

## MATERIALS and METHODS

### Experimental model and study participant details

The study cohort for Visium 10X and GeoMx DSP spatial transcriptomics analysis comprised 2 patients who initially received a diagnosis of IDH-mut WHO grade 2 astrocytoma and were followed throughout the course of their disease. Tumour samples were collected during initial and recurrent surgical resections at St. Michael’s Hospital (Unity Health Toronto, Toronto, Canada) carried out in accordance with approved guidelines and with patient consent under ethics approval REB #19-311. The clinical characteristics of the patient cohort are detailed in Table S1.

### Sample preparation

FFPE tissue sections for GeoMx DSP experiments were cut to a thickness of 5μm using a microtome and mounted onto Superfrost Plus slides within 1 month of spatial transcriptomics analysis. Slides were stored at 4°C with a desiccator and shipped to OICR for processing on GeoMx DSP (MH).

The RNA quality of each sample was evaluated by 2100 Agilent Bioanalyzer System after isolating RNA using the Qiagen RNeasy FFPE Kit (73504) from 30µm FFPE tissue scrolls and 10μm each for fresh-frozen tissue. Samples with DV200 values > 30% and an RNA integrity value > 7 were profiled by spatial transcriptomics.

### Histopathology

The adjacent sections of GeoMx slides and same sections of Visium slides were stained with haematoxylin and eosin (H&E) for histological examination.

Visium brightfield images were acquired at 10X magnification using a Zeiss Axio Imager 2 microscope.

GeoMx H&E slides were scanned using 3DHISTECH Pannoramic 250 digital slide scanner equipped with a Plan-Apochromat 20× objective and a CIS VCC-FC60FR19CL camera and processed using PANNORAMIC Viewer software (1.15.4).

The review of the histology slides and the selection of the target areas were performed by pathologist DM and AI. The area of interest was meticulously chosen focusing solely on the infiltrating-solid tumour boundary regions.

### GeoMx experimental procedure

The FFPE slides were baked for 2.5 hours, deparaffinized and rehydrated following the standard operating procedure for slide preparation of RNA assays outlined in MAN-10150-04. The slides were subjected to heat-induced epitope retrieval (1X Tris-EDTA pH 9.0 for 20 minutes at 100°C) followed by proteinase K digestion (0.1 μg/ml for 15 minutes at 37 °C). Following digestion tissue morphology was fixed in 10% neutral buffered formalin (NBF). The tissue sections were then hybridized with the Human Whole Transcriptome Atlas (WTA) probes (Bruker) overnight at 37°C. The following day sections underwent 2 x 25 min stringent washes to remove off target probes (1:1 4x SSC buffer & formamide), the slides were blocked and then incubated with morphology marker antibodies: GFAP (Texas Red, GA-5, 0.81 mg/ml), CD45 (Cy3, 2B11+PD7/26, 0.91 mg/ml), and Ki67 (Cy5, D3B5, 100 ug/mL). SYTO 13 provided in Bruker’s slide staining prep kit (FITC, 500 nM) was used as a nuclear stain.

Tissue sections were loaded into the GeoMx® Digital Spatial Profiling (DSP) instrument, which is a histology-based, spatial imaging platform where discrete regions of interest (ROIs) are selected for collection of oligo barcodes attached via an ultraviolet photocleavable linker to the in situ hybridization probes.

We selected ROIs for spatial analysis based on histological features that distinguished areas of infiltrative margins (“tumour edge”). Immunostaining for GFAP and Ki67 protein expression enabled the isolation of ROIs representing tumour cells (GFAP+) and highly proliferating tumour cells (GFAP+Ki67+).

UV light was then directed by the GeoMx® at each sample and released the RNA ID and UMI-containing oligonucleotide tags from the WTA probes for collection and sequencing preparation. Illumina i5 and i7 dual indexing primers were ligated to the oligonucleotide tags via PCR amplification to uniquely index each sample. AMPure XP beads (Beckman Coulter) were used for library purification. The final sequencing pool was quantified by qPCR and quality was assessed using the Bioanalyzer (Agilent). Sequencing was performed on an Illumina NovaSeq 6000 and fastq files were processed into gene count data for each sample using the GeoMx® NGS Pipeline. The sequencing run exceeded 90% sequencing saturation per ROI, indicating representative sampling.

### GeoMx data pre-processing

All computational analysis methods were run using defaults unless otherwise specified. Raw counts were grouped into a single dataset. The expression threshold used for this study is LOQ (limit of quantitation) = 2 and frequency of 10%. With these settings all targets with LOQ of 2 or greater in at least 10% of segments were kept. Filtering by LOQ gives confidence that the target is expressed above background levels and that it is accurately quantified.

Remaining genes were normalized by Q3 normalization.

### Dimensionality reduction and clustering

Principal Component Analysis (PCA) was performed on the normalized dataset using 21 principal components (ScanPy v.1.11; sc.pp.pca). To capture the local neighborhood structure of the data, a k-nearest neighbors graph was constructed with 9 neighbors (sc.pp.neighbors). Uniform Manifold Approximation and Projection (UMAP) was then applied for visualization of the high-dimensional data (sc.tl.umap). Clustering of cells was conducted using the Leiden algorithm with a resolution parameter set to 2, allowing identification of distinct cell populations (sc.tl.leiden). Cluster assignments were visualized on the UMAP embedding (sc.pl.umap).

### DEG analysis

Differentially expressed genes (DEGs) between Leiden clusters were identified using the Wilcoxon rank-sum test, selecting the top 90 genes per cluster (sc.tl.rank_genes_groups). DEGs were further filtered to retain genes with a minimum fold change of 1.5 and expressed in at least 25% of cells within the cluster. Differential gene expression between grade 4 samples and earlier progression stages was performed using DESeq2 (v1.34.0). Count data and sample metadata were input into a DESeqDataSet with a single-factor design comparing grade 4 versus other stages. DESeq2 normalization, dispersion estimation, and hypothesis testing were conducted with Cook’s distance-based outlier filtering and independent filtering applied. Genes with a mean normalized count ≥10, and absolute log2 fold change > 0.5 were considered differentially expressed. Because these preliminary analyses were generated from a limited number of patients and contain multiple spatial observations per specimen, we did not interpret adjusted p-values as evidence of grade-level statistical significance. Normalized counts were log-transformed and visualized using hierarchical clustering heatmaps.

### Gene set enrichment analysis

Gene set enrichment analysis was carried out using the enrichr function of GSEApy package. Reference gene sets used: GO_Biological_Process_2025, GO_Molecular_Function_2025, Reactome_Pathways_2024, KEGG. Enrichment was computed using the default parameters, terms with an adjusted p-value below 0.05 were considered.

### Correlation analysis between gene expression and proliferative index

Correlation between Ki67% and the expression of selected genes was quantified using scikit-learn LinearRegression model. Samples were grouped by grade. A linear regression model was fitted to the normalized gene expression to assess overall trends in expression-proliferation relationships across tumour progression stages.

### Visualization of gene expression

Expression patterns of marker genes were visualized using dot plots, violin plots, and matrix plots. Dot plots summarized gene expression across clusters with dendrograms illustrating cluster relationships (sc.pl.dotplot). Violin plots were generated for specific genes to compare expression distributions between clusters (sc.pl.violin). Matrix plots depicted scaled expression levels across developmental stages with hierarchical clustering (sc.pl.matrixplot).

### Visium experimental procedure

Spatial transcriptomics was performed using the 10X Genomics Visium CytAssist platform with the Human Transcriptome Probe Set v2.0. Fresh-frozen tissue sections (10 μm thickness) were mounted onto Visium CytAssist slides, with two samples per slide. Sample preparation— including H&E staining, imaging, probe hybridization, and library construction—was carried out following the manufacturer’s protocol (10X Genomics, Visium Spatial Protocols – Tissue Preparation Guide, CG000240, and Visium CytAssist Spatial Gene Expression for Fresh Frozen – Tissue Preparation Guide, CG000636, Rev A). Spatially barcoded probe hybridization was performed to capture mRNA, followed by reverse transcription and second-strand synthesis. Amplified cDNA libraries underwent dual SPRI size selection and were indexed with Illumina P5/P7 adapters. Library quality was assessed via Fragment Analyzer (Agilent) and qPCR quantification. Sequencing was conducted using an Illumina NovaSeq 6000 system with a 200-cycle v1.5 S4 flow cell, employing the standard 10x Visium GEX read configuration (28 bp for Read 1, 10 bp for i7 and i5 index reads, and 90–120 bp for Read 2). A minimum of 1% PhiX was used as a sequencing control. Sequencing targeted a depth of ∼50,000 reads per spot on the capture area, yielding a total of ∼98 million reads across 4 samples (equivalent to ∼25,000 reads/spot/sample on average). Preliminary data processing, including demultiplexing, alignment to the GRCh38 reference genome (version 2020-A), and UMI counting, was performed using the Space Ranger pipeline (v2.0.1) from 10x Genomics. Spatial gene expression matrices were generated using the Visium Human Transcriptome Probe Set v2.0.

### Visium data pre-processing and normalization

Spatial transcriptomics Visium data were processed using the Seurat (v4.3.0) and STUtility packages in R. Samples corresponding to histological grade 2, 3, and 4 gliomas were loaded and quality-filtered individually. Spots with high mitochondrial gene content (>3%) or elevated hemoglobin expression (>1%) were excluded to remove low-quality regions. Each grade-specific Seurat object was then normalized using the NormalizeData function (log-normalization with a scale factor of 10,000) and scaled (ScaleData). Filtered and pre-processed objects were then merged into a unified dataset using the merge() function, followed by JoinLayers() to integrate spatial layers.

### Feature selection and dimensionality reduction

Highly variable genes were identified using FindVariableFeatures(). For each individual sample, spatially variable features were further identified using Moran’s I statistic (nfeatures = 2000) on the “Spatial” assay. Principal component analysis was performed using the RunPCA() function to reduce data dimensionality, with the top 30 components retained based on elbow plot inspection. A shared nearest neighbor graph was constructed using FindNeighbors() (dims = 1:30), and clustering was performed using FindClusters() with a resolution of 0.4. UMAP embeddings were computed for visualization using RunUMAP(). Clusters with highest *MBP* expression (representing normal cells) were excluded from DEG analysis. Cellular and spatial domains were visualized with UMAP and spatial plots using DimPlot() and SpatialFeaturePlot() respectively.

### DEG analysis

Differential expression analysis was conducted using FindMarkers() for pairwise comparisons, and FindAllMarkers() to identify markers across all tumour grades. Marker gene expression was visualized using violin plots (VlnPlot) following filtering of zero-expression spots for each gene of interest. Spatially Variable genes (SVGs) were identified using SpatiallyVariableFeatures() function and visualized using SpatialFeaturePlot() in Seurat.

### Gene ontology and pathway enrichment analysis

Genes upregulated in grade 4 tumours (adjusted p-value < 0.05) were used for functional enrichment analysis. Gene Ontology enrichment for biological processes and molecular function was performed using the clusterProfiler::enrichGO() function with keyType = “SYMBOL” and OrgDb = org.Hs.eg.db. KEGG pathway enrichment analysis was also conducted using enrichKEGG() on genes mapped to Entrez IDs (bitr() conversion), using a p-value cutoff of 0.05. We did not interpret spot-level adjusted p-values as evidence of grade-level statistical significance.

### LASSO and EN regression models

RNA sequencing and clinical data for 446 patients were obtained from the TCGA Pan-Cancer Atlas^24^. Clinical metadata were extracted from the TCGA Clinical Data Resource (CDR), and gene expression data were downloaded from the TCGA RNASeqV2 dataset. IDH mutation status was annotated using a published reference dataset. Samples were filtered to include only IDH-mutant astrocytomas (grades 2–4).

A curated panel of genes implicated in glioma biology and spatial connectivity was selected from DEG analysis for survival modeling. Expression data from the merged TCGA dataset were filtered to retain samples with complete survival information (overall survival [OS] or progression-free interval [PFI] referred to as PFS). Expression values for selected genes of interest were transformed using log2(x + 1) prior to modeling.

The prognostic models were developed in the training set and evaluated in an independent held-out test set. The cohort was randomly split into 75% training and 25% testing subsets, separately for OS and PFS, as described by Raleigh et al.^26^ TCGA tumour grade was retained as an ordered factor and stored alongside the survival and gene-level predictors. Regularized models were trained exclusively in the training set, and the resulting risk scores were subsequently evaluated in the held-out test set.

Regularized Cox proportional hazards models were fit using the glmnet package (v4.1). LASSO models were fitted with α = 1 using 10-fold cross-validation. Elastic Net models were evaluated across α values ranging from 0.05 to 0.95, with the optimal α selected based on minimum mean cross-validated partial likelihood deviance. For each model, the penalty parameter λ was selected using `lambda.min`, corresponding to the value that minimized the mean cross-validated partial likelihood deviance. Non-zero coefficients were extracted to identify genes retained by each regularized model. Gene-derived risk scores were evaluated in the held-out test set using Cox proportional hazards models implemented in the survival package (v3.7-0), and model discrimination was quantified using the concordance index (C-index). Models incorporating TCGA tumour grade were additionally evaluated using multivariable Cox regression to determine whether the gene-derived risk scores provided prognostic information beyond tumour grade. Model fit was compared using likelihood-ratio tests. For exploratory Kaplan-Meier analyses, patients were stratified into high- and low-risk groups using an outcome-oriented cut-point determined with the `surv_cutpoint` function in survminer (v0.5.0), and survival distributions were compared using log-rank tests. All analyses were performed in R (v4.3.2).

### DNA extraction

DNA was extracted from FFPE Grade 4 tumour samples as per manufacturer’s protocol (Mag-Bind FFPE DNA/RNA 96 Kit (Omega, #M6955) & MagMAX DNA Multi-Sample Ultra 2.0 Kit (ThermoFisher A36570)) on the KingFisher™ Flex Purification System equipped with the 96 Deep-well Head (Thermofisher Cat#5400630).

### WES

100 ng of DNA were used for preparation of targeted sequencing library. DNA was sheared to a targeted average fragment size of 250 bp using a E220 Focused-Ultrasonicator (Covaris). Pre-capture libraries were prepared from the sheared DNA using KAPA DNA HyperPrep Kit (Roche Cat# 07962363001) with IDT xGen Duplex Seq Adapter – Tech Access (IDT Cat# 1080799). Target capture with the IDT xGEN Exome Research Panel v2 (IDT Cat# 10005153) was performed using IDT xGEN Hybridization and Wash Kit (IDT Cat# 1080584) and IDT xGEN Universal Blockers – TS mis (IDT Cat# 1075475) according to manufacturer’s instruction. Prepared libraries were balanced, pooled and loaded on an Illumina NovaSeq X Plus Sequencing System (Illumina Cat# 20084804) and sequenced at 2×151 cycles.

### Quantification and statistical analysis

The Wilcoxon rank-sum test was used to compare the means between two groups, such as gene expression, unless otherwise specified. All statistical tests in this study were by default two-sided tests. For multiple-testing correction, we applied the Benjamini–Hochberg’ method to compute the FDR. Results with p□values or FDR less than 0.05 were considered statistically significant. Significance levels were indicated as follows: *p□<□0.05, **p□<□0.01, ***p < 0.001, ****p□<□0.0001.

## Required Statements

### Ethics

St. Michael’s Hospital REB #19-311. The Hospital for Sick Children REB #1000024587.

### Authorship

Conception and design: AI, SD

Development of methodology: AI, SA, SD, MS, MH

Acquisition of data: AI, MH, FA

Analysis and interpretation of data: AI, SA

Writing, review, and/or revision of the manuscript: AI, SA, SD, DGM, RGWV, DPC, TJP, MW

Administrative, technical, or material support (i.e., reporting or organizing data, constructing databases): AI, MW, MH, FA, DGM

Study supervision: SA, SD

### Data Availability

The raw and processed spatial transcriptome data generated in this study have been deposited at the National Center for Biotechnology Information (NCBI) Gene Expression Omnibus (GEO) with accession number GSE318238. Raw WES data generated in this study have been deposited at NCBI with project ID PRJNA1460314 and is available for viewing at cBioportal (IDHMA_Das).

### Code availability

All R and Python scripts supporting the findings of this study are available in the GitHub repository at https://github.com/alyonaivanova22/Spatially-resolved-gene-expression-across-malignant-transformation-of-IDH-mut-glioma.git

## REFERENCES

1. Miller JJ, et al. Isocitrate dehydrogenase (IDH) mutant gliomas: a Society for Neuro-Oncology (SNO) consensus review on diagnosis, management, and future directions. Neuro Oncol. 2023;25(1):4–25. doi:10.1093/neuonc/noac207

2. Dang L, et al. Cancer-associated IDH1 mutations produce 2-hydroxyglutarate. Nature. 2009;462(7274):739–744. doi:10.1038/nature08617

3. Ward PS, et al. The common feature of leukemia-associated IDH1 and IDH2 mutations is a neomorphic enzyme activity converting α-ketoglutarate to 2-hydroxyglutarate. Cancer Cell. 2010;17(3):225–234. doi:10.1016/j.ccr.2010.01.020

4. Pusch S, et al. Mutant IDH1 R132H increases production of 2-hydroxyglutarate in glial cells in vitro and in vivo. Acta Neuropathol. 2014;128(3):341–353. doi:10.1007/s00401-014-1307-7

5. Wei Y, et al. Stalled oligodendrocyte differentiation in IDH-mutant gliomas. Genome Med. 2023;15:24. doi:10.1186/s13073-023-01175-6

6. Louis DN, et al. The 2021 WHO classification of tumours of the central nervous system: a summary. Neuro Oncol. 2021;23(8):1231–1251. doi:10.1093/neuonc/noab106

7. Xia L, et al. Prognostic role of IDH mutations in gliomas: a meta-analysis of 55 observational studies. Oncotarget. 2015;6:17354–17365.

8. Turkalp Z, et al. IDH mutation in glioma: new insights and promises for the future. JAMA Neurol. 2014;71(10):1319–1325. doi:10.1001/jamaneurol.2014.1205

9. Han S, et al. IDH mutation in glioma: molecular mechanisms and potential therapeutic targets. Br J Cancer. 2020;122(11):1580–1589. doi:10.1038/s41416-020-0814-x

10. Solomou G, et al. Mutant IDH in gliomas: role in cancer and treatment options. Cancers (Basel). 2023;15(11):2883. doi:10.3390/cancers15112883

11. Yan H, et al. IDH1 and IDH2 mutations in gliomas. N Engl J Med. 2009;360(8):765–773. doi:10.1056/NEJMoa0808710

12. Parsons DW, et al. An integrated genomic analysis of human glioblastoma multiforme. Science. 2008;321(5897):1807–1812. doi:10.1126/science.1164382

13. Giese A, et al. Cost of migration: invasion of malignant gliomas and implications for treatment. J Clin Oncol. 2003;21(8):1624–1636. doi:10.1200/JCO.2003.05.063

14. Hatzikirou H, et al. “Go or grow”: the key to the emergence of invasion in tumour progression? Math Med Biol. 2012;29(1):49–65. doi:10.1093/imammb/dqq011

15. Seifert M, Schackert G, Temme A, et al. Molecular Characterization of Astrocytoma Progression Towards Secondary Glioblastomas Utilizing Patient-Matched Tumor Pairs. Cancers (Basel). 2020;12(6):1696. Published 2020 Jun 26. doi:10.3390/cancers120616

16. Venkatesh HS, et al. Electrical and synaptic integration of glioma into neural circuits. Nature. 2019;573(7775):539–545. doi:10.1038/s41586-019-1563-y

17. Venkatesh HS, et al. Neuronal activity promotes glioma growth through neuroligin-3 secretion. Cell. 2015;161(4):803–816. doi:10.1016/j.cell.2015.04.012

18. Venkataramani V, et al. Synaptic input to brain tumours: clinical implications. Neuro Oncol. 2021;23(1):23–33. doi:10.1093/neuonc/noaa158

19. Hsieh AL, et al. Widespread neuroanatomical integration and distinct electrophysiological properties of glioma-innervating neurons. Proc Natl Acad Sci U S A. 2024;121(50):e2417420121. doi:10.1073/pnas.2417420121

20. Drexler R, et al. Epigenetic neural glioblastoma enhances synaptic integration and predicts therapeutic vulnerability. bioRxiv. Preprint. Posted August 7, 2023. doi:10.1101/2023.08.04.552017

21. Drummond KJ, et al. Perioperative IDH inhibition in treatment-naive IDH-mutant glioma: a pilot trial. Nat Med. 2025;31:3451–3463. doi:10.1038/s41591-025-03884-4

22. Taylor KR, et al. Glioma synapses recruit mechanisms of adaptive plasticity. Nature. 2023;623:366–374. doi:10.1038/s41586-023-06678-1

23. Mortazavi A, et al. IDH-mutated gliomas promote epileptogenesis through d-2-hydroxyglutarate-dependent mTOR hyperactivation. Neuro Oncol. 2022;24(9):1423–1435. doi:10.1093/neuonc/noac003

24. Cancer Genome Atlas Research Network. The Cancer Genome Atlas Pan-Cancer analysis project. Nat Genet. 2013;45(10):1113–1120. doi:10.1038/ng.2764

25. Greenwald AC, et al. Integrative spatial analysis reveals a multi-layered organization of glioblastoma. Cell. 2024;187(10):2485–2501.e26. doi:10.1016/j.cell.2024.03.029

26. Raleigh D, et al. Spatial synaptic connectivity underlies oligodendroglioma evolution and recurrence. Research Square. Preprint. Posted April 4, 2025. doi:10.21203/rs.3.rs-6299872/v1

27. Yan L, et al. Benchmarking and integration of methods for deconvoluting spatial transcriptomic data. Bioinformatics. 2023;39(1):btac805. doi:10.1093/bioinformatics/btac805

28. Saqib J, et al. From pixels to cell types: a comprehensive review of computational methods for spatial transcriptomics deconvolution. Genom Inform. 2025;23:22. doi:10.1186/s44342-025-00055-2

29. Hu J, et al. Statistical and machine learning methods for spatially resolved transcriptomics with histology. Comput Struct Biotechnol J. 2021;19:3829–3841. doi:10.1016/j.csbj.2021.06.052

30. Johnson BE, Mazor T, Hong C, et al. Mutational analysis reveals the origin and therapy-driven evolution of recurrent glioma. Science. 2014;343(6167):189–193.

31. Sottoriva A, Spiteri I, Piccirillo SGM, et al. Intratumor heterogeneity in human glioblastoma reflects cancer evolutionary dynamics. Proc Natl Acad Sci U S A. 2013;110(10):4009–4014.

32. Nicholson JG, Fine HA. Diffuse glioma heterogeneity and its therapeutic implications. Cancer Discov. 2021;11(3):575–590.

33. Campbell BB, Light N, Fabrizio D, et al. Comprehensive analysis of hypermutation in human cancer. Cell. 2017;171(5):1042–1056.

34. Rayner E, van Gool IC, Palles C, et al. A panoply of errors: polymerase proofreading domain mutations in cancer. Nat Rev Cancer. 2016;16(2):71–81.

35. Le DT, Durham JN, Smith KN, et al. Mismatch repair deficiency predicts response of solid tumors to PD-1 blockade. Science. 2017;357(6349):409–413.

36. Cancer Genome Atlas Research Network, Brat DJ, Verhaak RG, et al. Comprehensive, Integrative Genomic Analysis of Diffuse Lower-Grade Gliomas. N Engl J Med. 2015;372(26):2481–2498. doi:10.1056/NEJMoa1402121

37. Donehower LA, Soussi T, Korkut A, et al. Integrated analysis of TP53 gene and pathway alterations in cancer. Cell Rep. 2019;28(5):1370–1384.e5.

38. Mantovani F, Collavin L, Del Sal G. Mutant p53 as a guardian of the cancer cell. Nat Rev Cancer. 2019;19(2):101–114.

39. Kumar MS, Lu J, Mercer KL, et al. Impaired microRNA processing enhances cellular transformation and tumorigenesis. Nat Genet. 2007;39(5):673–677.

40. Appin CL, Brat DJ. Molecular genetics of gliomas. Cancer J. 2014;20(1):66–72.

41. Eckel-Passow JE, Lachance DH, Molinaro AM, et al. Glioma groups based on 1p/19q, IDH, and TERT promoter status. N Engl J Med. 2015;372(26):2499–2508.

42. Abounader R, Laterra J. Scatter factor/hepatocyte growth factor in brain tumor growth and angiogenesis. Neuro Oncol. 2005;7(4):436–451.

43. Verhaak RGW, Hoadley KA, Purdom E, et al. Integrated genomic analysis identifies clinically relevant glioblastoma subtypes. Cancer Cell. 2010;17(1):98–110.

44. Fruman DA, Chiu H, Hopkins BD, et al. The PI3K pathway in human disease. Cell. 2017;170(4):605–635

45. Arita H, Narita Y, Fukushima S, et al. Upregulating mutations in the TERT promoter commonly occur in adult malignant gliomas. Acta Neuropathol. 2013;126(2):267–276.

46. Labussière M, Boisselier B, Mokhtari K, et al. Combined analysis of TERT, IDH, and 1p/19q status defines glioma subsets. Neurology. 2014;83(14):1200–1206.

